# NEURAL POPULATION DYNAMICS IN MOTOR CORTEX ARE DIFFERENT FOR REACH AND GRASP

**DOI:** 10.1101/667196

**Authors:** Aneesha K. Suresh, James M. Goodman, Elizaveta V. Okorokova, Matthew T. Kaufman, Nicholas G. Hatsopoulos, Sliman J. Bensmaia

## Abstract

Rotational dynamics are observed in neuronal population activity in primary motor cortex (M1) when monkeys make reaching movements. This population-level behavior is consistent with a role for M1 as an autonomous pattern generator that drives muscles to produce movement. Here, we show that M1 does not exhibit smooth dynamics during grasping movements, suggesting a more input-driven circuit.

## MAIN TEXT

Populations of neurons in primary motor cortex (M1) exhibit smooth dynamics in their responses when animals make reaching or cycling movements ^1–4^. One interpretation of this population-level behavior is that M1 acts a pattern generator that drives muscles to give rise to movement. A major question is whether smooth population dynamics reflect a general principle of M1 function, or whether they underlie some behaviors but not others. To address this question, we examined the degree to which M1 exhibits smooth rotational dynamics during grasping movements, which involve a plant with a different function, more joints, and different mechanical properties than the arm, and which is subserved by a different portion of M1.

To this end, we recorded the neural activity in M1 and somatosensory cortex (SCx) using chronically implanted electrode arrays as monkeys performed a grasping task, restricting our analysis to responses before object contact (Figure S1). Animals were required to hold their arms still at the elbow and shoulder joints as a robotic arm presented each object to their contralateral hand. This task limits proximal limb movements and isolates grasping movements. For comparison, we also examined the responses of M1 populations during a center-out reaching task ^5^.

First, we characterized the population dynamics in M1 during reaching and grasping movements (Figure 1). We used jPCA to search for rotational dynamics in a low-dimensional manifold of M1 population activity ^1^. Replicating previous findings, reaching evoked multiphasic activity in single M1 neurons (Figure 1A) and strong rotational dynamics in the population (Figure 1C). During grasp, individual M1 neurons again exhibited strong, multiphasic modulation (Figure 1B), but rotational dynamics were weak or absent (Figure 1D,E).

**Figure 1.**
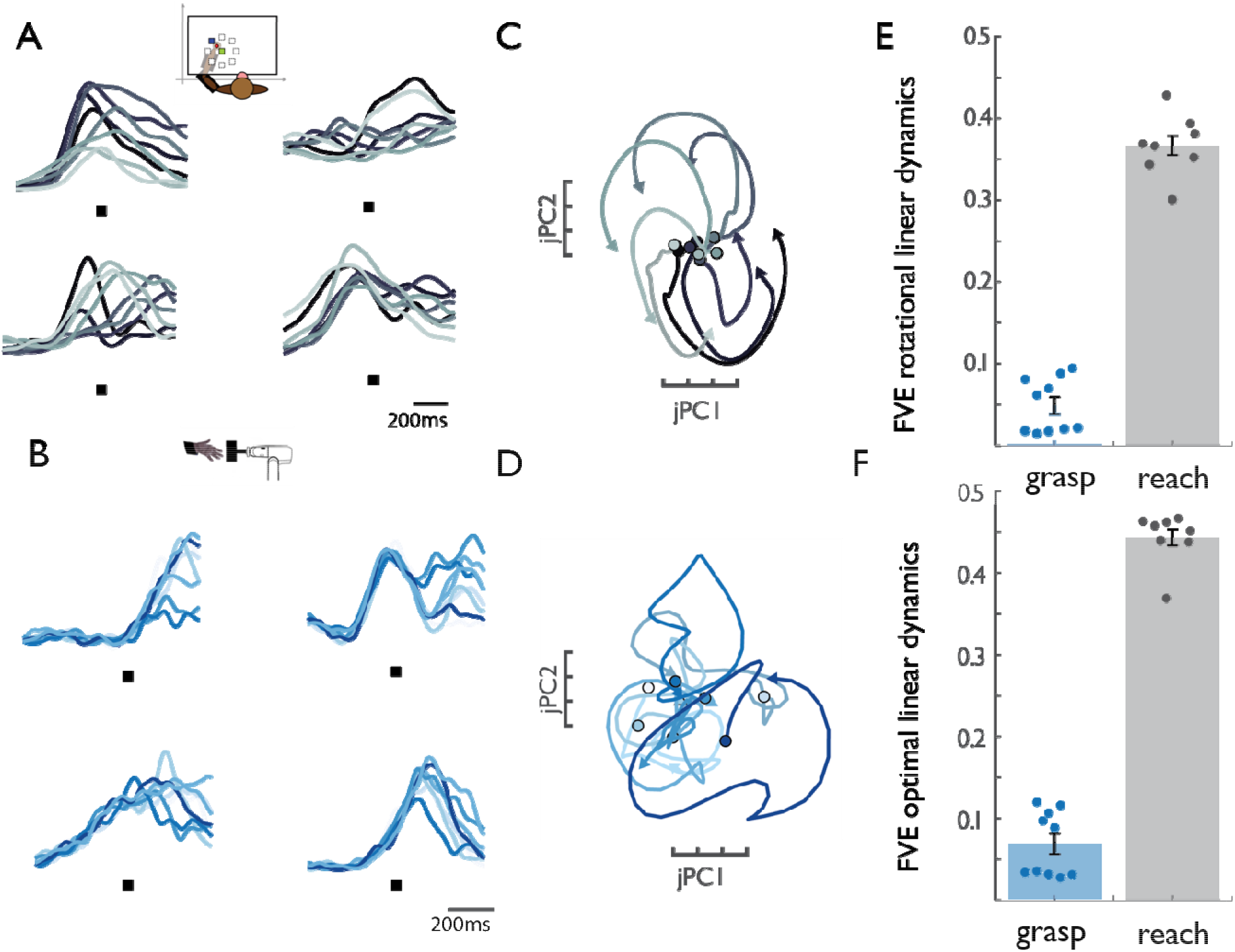
M1 rotational dynamics during reaching and grasping. **A|** Normalized peri-event histograms aligned to movement onset (black square) for 4 representative neurons during the reaching task. Each shade of gray indicates a different reach direction. **B|** Normalized peri-event histograms aligned to maximum aperture (black square) for 4 representative neurons during the grasping task. Each shade of blue indicates a different object group (see supplementary materials). **C|** Rotational dynamics in the population response during reaching along the first jPCA plane. Different shades of gray denote different reach directions. **D|** M1 rotational dynamics during grasping. Different shades of blue indicate different object groups. **E|** FVE (fraction of variance explained) in the rate of change of neural PCs (d*x*/dt) explained by the optimal rotational dynamical system. Difference is significant (two-sample two-sided equal-variance *t*-test, t(16) = −19.44, *p*=4.67e-13). Error bars denote standard error of the mean, and data points represent cross-validated results for 2 monkeys. **F|** FVE in the rate of change of neural PCs (d*x*/dt) explained by the optimal linear dynamical system. Difference is significant (two-sample two-sided equal-variance *t*-test, t(16) = −21.37 *p*=1.57e-14). Error bars denote standard error of the mean, and data points represent cross-validated results for 2 monkeys.

Given the poor fit of rotational dynamics to neural activity during grasp, we assessed whether activity could be described by a linear dynamical system of any kind. To test for linear dynamics, we fit a regression model using the first 10 principal components of the M1 population activity (*x*(t)) to predict their rates of change (d*x*/dt). We found *x*(t) to be far less predictive of d*x*/dt in grasp than in reach, suggesting much weaker linear dynamics in the neural representation of grasp than reach (Figure 1F). We verified that these results were not an artifact of data alignment, analysis interval, peak firing rate, or population size (Figure S2).

The possibility remains that dynamics are present in M1 during grasp, but that they are higher-dimensional than during reach, or that they are nonlinear. Indeed, previous work analyzing neural state spaces in M1 during a reach-grasp-manipulate task found that neural activity is higher-dimensional than that observed during reach movements alone ^6^. As a first test of these possibilities, we examined the relationship between movement and neural activity from the standpoint of decoding. We used a powerful recent technique, latent factor analysis via dynamical systems (LFADS), which infers and exploits latent dynamics to improve estimation of single-trial firing rates. Naturally, this benefit is only realized if the neural population acts like a dynamical system. Importantly, such dynamics are minimally constrained and can, in principle, be arbitrarily high dimensional and/or highly nonlinear. We then used a standard Kalman filter to decode joint angle kinematics from the inferred latent factors (Figure 2). If latent dynamics are present in the system, LFADS should substantially improve kinematic decoding relative to simple Gaussian-smoothed spike trains. Replicating previous results, decoding accuracy was substantially improved for reaching when inferring firing rates using LFADS (Figure 2A,C). However, LFADS offered no accuracy improvement when decoding grasping kinematics (Figure 2B,C), even though grasp was decoded just as well as reach under Gaussian smoothing of spike trains. These effects were consistent across all 30 degrees of freedom of the hand (Figure S3). These decoding results demonstrate that the strong dynamical structure seen in M1 population activity during reaching is not observed during grasp, even when dimensionality and linearity constraints are lifted.

**Figure 2.**
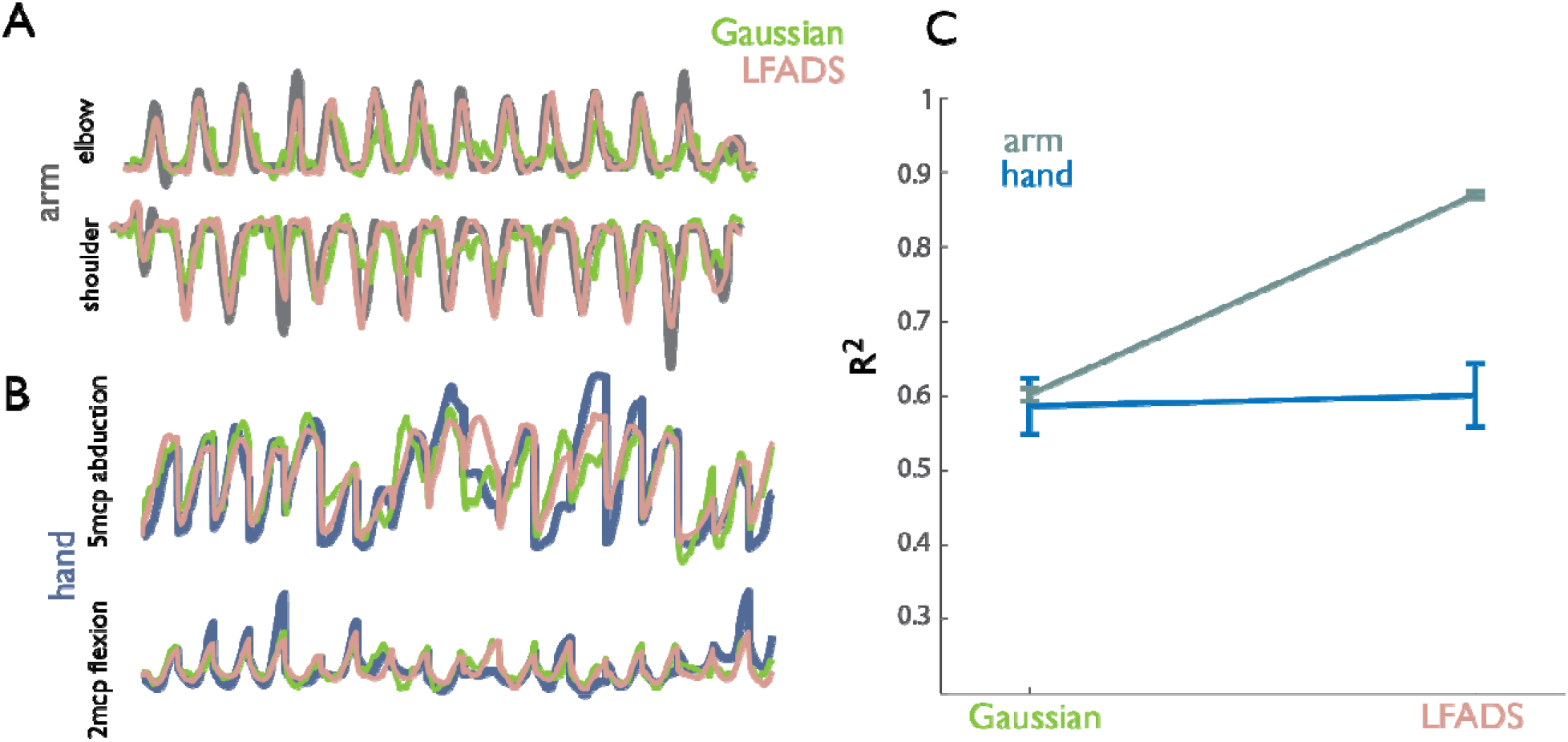
Accuracy of Kalman filter decoders of kinematics using neural data pre-processed with Gaussian smoothing or with the assumption of underlying latent dynamics (LFADS). **A,B|** Example kinematic traces reconstructed with and without the assumption of dynamics. **A|** Angles of arm joints (gray) along with angles decoded when neuronal responses are preprocessed with Gaussian kernel (green) and with LFADS (pink). **B|** Angles of hand joints (blue) along with their decoded counterparts (Gaussian smooth in green, LFADS in pink). **C|** Mean performance of decoders for the arm (2 DoF) and the hand (22 DoF), 10-fold cross-validated using a population of 44 neurons. Individual joint data shown in Supplementary Figure 3. LFADS leads to substantial improvements over typical pre-processing of neural data (Gaussian smoothing) for decoding reaching but not hand kinematics.

As a separate way to gauge the presence of nonlinear dynamics in grasping responses, we computed a neural ‘tangling’ metric, which assesses the degree to which network dynamics are governed by a smooth and consistent flow field ^3^. In a smooth, autonomous dynamical system, neural trajectories passing through nearby points in state space have similar derivatives. The tangling metric (*Q*) assesses the degree to which this is the case over a specified (reduced) number of dimensions (Figure S4). During reaching, muscle activity and movement kinematics have been shown to exhibit more tangling than does M1 activity, presumably because the neural system acts as a pattern generator while muscles are input-driven ^3^. We replicated these results for reaching: neural activity was much less tangled than the corresponding arm kinematics (position, velocity, and acceleration of joint angles)(Figure 3A). For grasp, however, M1 activity was somewhat *more* tangled than were the corresponding hand kinematics (Figure 3B). Next, we compared tangling in M1 to tangling in SCx, which, as a sensory area, is expected to exhibit tangled activity ^3^. Surprisingly, M1 and SCx activity was similarly tangled during grasp (Figure 3C). In summary, then, M1 responses during grasp do not exhibit the properties of an autonomous dynamical system but rather resemble sensory responses (Figure 3D).

**Figure 3.**
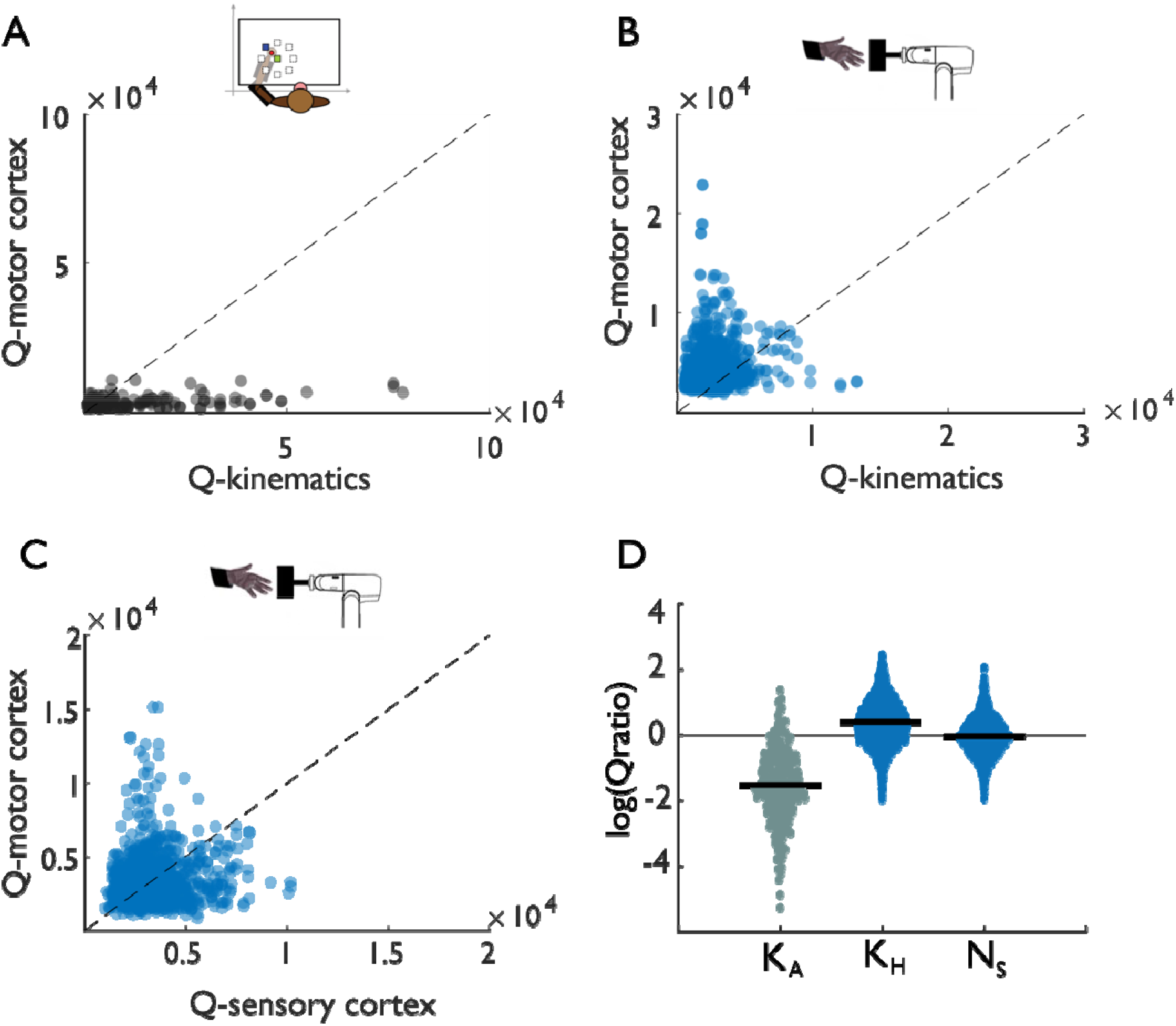
Tangling in reach and grasp. **A|** Tangling metric (Q) for population responses in motor cortex during reaching vs. Q for reaching kinematics. Kinematic tangling is higher than neural tangling, consistent with motor cortex acting as a pattern generation during reach. **B|** Q-M1 population vs. Q-kinematics for grasping. Neural tangling is *higher* than kinematic tangling, which argues against pattern generation as the dominant drive during grasp. **C|** Q-M1 population vs. Q-SCx population. Neural tangling is similar in M1 and SCx, suggesting that M1 is as input driven as a sensory area. **D|** Summary results for two monkeys: Log of Q-motor/Q-kinematics of the arm during reach (K_A_), Q-motor/Q-kinematics of the hand during grasp (K_H_), and Q-motor/Q-sensory during grasp (N_s_). Black bars denote mean. Differences are significant to reaching (two-sample two-sided equal-variance *t*-test: K_H_ | t(2978)=−43, *p*=1.03e-130; N_s_ |t(2978)=−39 *p*=1.87e-121).

We speculate that the similar level of tangling observed in SCx and M1 may reflect greater interplay between these areas during grasp than during proximal limb movements. Indeed, sensory feedback can obscure the intrinsic dynamics in neuronal circuits ^7^ and sensory representations are highly tangled ^3^. Additionally, the relatively greater importance of sensory feedback during grasp as opposed to reach is well-documented: monkeys can perform pointing movements with reasonable accuracy even in the absence of sensory input ^8^, whereas substantial deficits in finger coordination during grasp are observed when inactivating SCx ^9^. Furthermore, orderly dynamics have been observed during a single-finger pointing task in human subjects with ALS ^10^, a condition wherein neuronal loss characterizes both M1 and SCx ^11^. That orderly M1 dynamics during hand and finger movements emerge after such loss is broadly consistent with SCx-M1 communication playing a role in tangling the M1 activity. Moreover, quick reaching movements are associated with a stereotyped triphasic muscle activation pattern ^12^ that is observed regardless of sensory feedback. To our knowledge, similar patterns of muscle activity have not been documented during grasping movements. Finally, projections between primate somatosensory and motor cortices support communication between the two ^13–15^, demonstrating a pathway by which M1 and SCx dynamics might be directly linked rather than driven by parallel inputs. While this pathway is not unique to the hand, it may be more engaged during grasp than reach.

In conclusion, we show that the lawful dynamics observed in M1 during reaching are not observed during grasping, and propose that this difference may be due to increased interplay between somatosensory and motor cortices during grasp.

## METHODS

### Behavior and neurophysiology for grasping task

We recorded single- and multi-unit responses in both the primary motor and somatosensory cortices (M1 and SCx) of two monkeys (Macaca mulatta) (M1: N_1_ = 53, N_2_ = 58 | SCx: N_1_ = 28 N_2_ = 26), and from M1 of a third monkey (M1: N_3_ = 80), as they grasped each of 35 different objects an average of 10 times. We only used neural responses from Monkey 3 in the decoding analysis (from M1 responses) because we did not obtain enough data from this animal’s SCx. Neural recordings were obtained using semi-chronic electrode arrays (SC96 arrays, Gray Matter Research, Bozeman, MT) ^16^ across six and nine sessions for Monkeys 1 and 2, respectively. Electrodes, which were individually depth-adjustable, were moved to different depths on different sessions to capture new units. Units from Monkey 3 were recorded across two sessions from Utah electrode arrays (UEAs, Blackrock Microsystems, Inc., Salt Lake City, UT) and floating microelectrode arrays (FMAs, Microprobes for Life Science, Gaithersburg, MD) targeting rostral and caudal subdivisions of the hand representation of M1, respectively. Single units from all sessions (treated as distinct units) were extracted manually using an Offline Sorter (Plexon Inc., Dallas TX). Units were identified based on inter-spike interval distribution and waveform shape and size.

Hand joint kinematics, namely the angles and angular velocities about all motile axes of rotation in the joints of the wrist and digits, were tracked at a rate of 100 Hz by means of a 14-camera motion tracking system (MX-T series, VICON, Los Angeles, CA). The VICON system tracked the three-dimensional positions of the markers, and joint angle kinematics were computed using inverse kinematics based on a musculoskeletal model of the human arm (https://simtk.org/projects/ulb_project) ^17–23^ implemented in Opensim (https://simtk.org/frs/index.php?group_id=91) ^24^ with segments scaled to the sizes of those in a monkey limb. Task and kinematic recording methods are similar to previously reported ones ^25^, but with a greater number of objects (35) and more detailed kinematic reconstructions (30 joints) ^26^.

All surgical, behavioral, and experimental procedures conformed to the guidelines of the National Institutes of Health and were approved by the University of Chicago Institutional Animal Care and Use Committee.

### Behavior and neurophysiology for reaching task

To compare grasp to reach, we analyzed previously-published single- and multi-unit responses from M1 of two additional monkeys (Macaca mulatta) (M1: N_4_ = 76,, N_5_ = 107) operantly trained to move a cursor in a variable-delay center-out reaching task ^5^. The monkey’s arm rested on cushioned arm troughs secured to links of a two-joint exoskeletal robotic arm (KINARM system; BKIN Technologies, Kingston, Ontario, Canada) underneath a projection surface. The shoulder and elbow joint angles were sampled at 500 Hz by the motor encoders of the robotic arm, and the x and y positions of the hand were computed using the forward kinematic equations. The center-out task involved movements from a center target to one of eight peripherally positioned targets (5 to 7 cm away). Targets were radially defined, spanning a full 360 degree rotation about the central target in 45 degree increments. Each trial comprised two epochs: first, an instruction period lasting 1 to 1.5 s, during which the monkey held its hand over the center target to make the peripheral target appear; second, a “go” period, cued by blinking of the peripheral target, which indicated to the monkey that it could begin to move toward the target. Single- and multi-unit activity from each monkey was recorded from a UEA implanted into the upper limb representation of contralateral M1.

All surgical, behavioral, and experimental procedures conformed to the guidelines of the National Institutes of Health and were approved by the University of Chicago Institutional Animal Care and Use Committee.

### Data Analysis

#### Rotational Dynamics

##### Data pre-processing

For both the reach and grasp datasets, neuronal responses were aligned to the start of movement, resampled at 100 Hz so that reach and grasp data were at the same time resolution, averaged across trials, then smoothed by convolution with a Gaussian (20 ms S.D.). We then followed the same data pre-processing steps as outlined in Churchland et al. 2012: normalization of individual neuronal firing rates, subtraction of the cross-condition mean peri-event time histogram (PETH) from each neuron’s response in each condition, and applying principal component analysis (PCA) to reduce the dimensionality of the population response. We used 10 dimensions instead of six (cf. Churchland et al. 2012) as a compromise between the lower-dimensional reach data and the higher-dimensional grasp data.

##### jPCA

We then applied to the population data (reduced to 10 dimensions by PCA) a published dimensionality reduction method, jPCA ^1^, which finds orthonormal basis projections that capture rotational structure in the data. Specifically, the neural state is compared with its derivative and the strictly rotational dynamical system that explains the largest fraction of variance in that derivative is identified. The delay periods between the presentation/go-cue for the datasets varied, along with the reaction times, so we analyzed over time intervals (~500 ms) that maximized rotational variance for each dataset. For the reach data, data were aligned to the start of movement and the analysis window was centered on this event, whereas for the grasp data, data were aligned to maximum hand aperture, and we analyzed the interval centered on this event. In some cases, the center of this 500-ms window was shifted between −250 ms to +250 ms relative to the alignment event to obtain an estimate of how rotational dynamics change over the course of the trial (e.g., Figure S2). These events were chosen for alignment as they were associated with both the largest peak firing rates and the strongest rotational dynamics. Other alignment events were also tested, to validate robustness (Figure S2B).

##### Object clustering

Each of the 35 objects was presented 10 times per session, which yields a smaller number of trials per condition than were used to assess jPCA during reaching (at least 40). To permit pooling across a larger number of trials when visualizing and quantifying population dynamics with jPCA (Figure 1), objects in the grasp task were grouped into eight object clusters on the basis of the trial-averaged similarity of hand posture across all 30 joint degrees of freedom 10 ms prior to grasp (i.e., object contact). Objects were hierarchically clustered into 8 clusters on the basis of the Ward linkage function (MATLAB `clusterdata`). Eight clusters were chosen to match the number of conditions in the reaching dataset. Cluster sizes were not uniform; the smallest comprised 2 and the largest 9 different objects, with the median cluster comprising 4 objects.

As the clustering method just described yielded different cluster sizes, we assessed an alternative clustering procedure (Figure S2E) that guaranteed objects were divided into 7 equally-sized clusters (5 objects per cluster). Rather than determining cluster membership on the basis of a linkage threshold, cluster linkages were instead used to sort the objects on the basis of their dendrogram placements (MATLAB `dendrogram`). Clusters were obtained by grouping the first five objects in this sorted list into a common cluster, then the next five, and so on. This resulted in slightly poorer performance of jPCA (see *Quantification*).

For completeness, we also assess jPCA without clustering (Figure S2E). This also resulted in slightly poorer performance of jPCA and, by virtue of comprising 35 distinct neural trajectories instead of just 8, was considerably more difficult to visualize.

##### Quantification

In a linear dynamical system, the derivative of the state is a linear function of the state. We wished to assess whether a linear dynamical system could closely describe the neural activity. To this end, we first produced a denoised low-dimensional neural state (*X*) by reducing the dimensionality of the neuronal responses to 10 using PCA. Second, we numerically differentiated *X* to produce the empirical derivative, 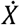. Next, we used regression to fit a linear model, predicting the derivative of the neuronal state from the current state: 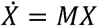. Finally, we computed the fraction of variance explained (FVE) by this model:

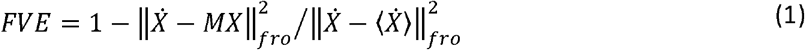

*M* was constrained to be skew-symmetric (*M*_*skew*_) unless otherwise specified; 〈⋅〉 indicates the mean of a matrix across samples, but not across dimensions; and ∥⋅∥_*fro*_ indicates the Frobenius norm of a matrix. Reaching data was 4-fold cross-validated, while grasp data was 5-fold cross-validated.

#### Control comparisons between arm and hand data

We performed several controls comparing arm and hand data to ensure that our results were not an artifact of trivial differences in the data or pre-processing steps.

First, we considered whether alignment of the data to different events might impact results. For the arm data, we aligned each trial to target onset and movement onset (Figure S2A). For the hand data, we aligned each trial to presentation of the object, movement onset, and the time at which the hand reached maximum aperture during grasp (Figure S2B). Rotational dynamics were strongest (though still very weak) when neuronal responses were aligned to maximum aperture so this alignment is reported throughout the main text.

Second, we assessed whether rotations might be obscured due to differences in firing rates in the hand vs. arm responses. To this end, we compared peak firing rates for trial-averaged data from the arm and hand after pre-processing (excluding normalization) to directly contrast the inputs to the jPCA analysis given the two effectors/tasks (Figure S2C). Peak firing rates were actually higher for the hand than the arm, eliminating the possibility that our failure to observe dynamics during grasp was an artifact of weak responses.

Finally, we assessed whether differences in the sample size might contribute to differences in variance explained. To this end, we took five random samples of 55 neurons from the reaching data set – chosen to match the minimum number of neurons in the grasping datasets – and computed the cross-validated fraction of variance explained by the rotational dynamics. The smaller samples yielded identical fits as the full sample.

##### Tangling

We computed tangling of the neural population data (reduced to 15 dimensions by PCA) using a published method ^3^. In brief, the tangling metric estimates the extent to which neural population trajectories are inconsistent with what would be expected if they were governed by an autonomous dynamical system, with smaller values indicating consistency with such dynamical structure. Specifically, tangling measures the degree to which similar neural states, either during different movements or at different times for the same movement, are associated with different derivatives. This is done by finding, for each neural state (indexed by *t*), the maximum value of the tangling metric *Q*(*t*) across all other neural states (indexed by *t*’):

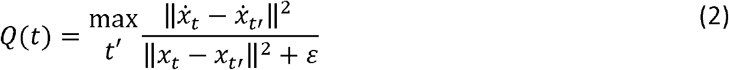

Here, *x*_*t*_ is the neural state at time *t* (a 15 dimensional vector containing the neural responses at that time), 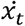 is the temporal derivative of the neural state, and ||⋅|| is the Euclidean norm, while *ε* is a small constant added for robustness to noise ^3^. This analysis is not constrained to work solely for neural data; indeed, we also apply this same analysis to trajectories of joint angular kinematics to compare with the tangling of neural trajectories.

The neural data were pre-processed using the same alignment, trial averaging, smoothing, and normalization methods described above. Joint angles were collected for both hand and arm data. For this analysis, joint angle velocity and acceleration were computed (six total dimensions for arm, 90 dimensions for hand). For reaching, we analyzed the epoch from 200 ms before to 100 ms after movement onset. For grasping, we analyzed the epoch starting 200 ms before to 100 ms after maximum aperture. Neuronal responses were binned in 10 ms bins to match the sampling rate of the kinematics.

We tested tangling at different dimensionalities and selected the dimensionality at which Q had largely leveled off for both the population neural activity and kinematics (Figure S4), namely 6 for reach kinematics (the maximum) and 15 for grasp kinematics and for the neuronal responses.

#### Decoding

We analyzed the extent to which we can decode kinematics of the hand and the arm using neural population activity recorded from primary motor cortex and compared performance with and without the assumption of underlying dynamics. To this end, we used the responses from monkey 3 (2 sessions with 44 and 36 M1 neurons and 20 kinematic DoF) performing the grasping task, and monkey 4 (64 M1 neurons and 2 kinematic DoF) performing the reaching task. We analyzed 800 ms of neural data preceding maximum aperture of the hand in the grasping task, and neural data from 600 ms before to 200 ms after movement onset in the reaching task.

##### Preprocessing

For decoding, we preprocessed the neural data using one of the two methods: smoothing with a Gaussian kernel (σ = 20 ms) or latent factor analysis via dynamical systems (LFADS, Pandarinath et al., 2018).

LFADS is a generative model that assumes that observed spiking arises from an underlying dynamical system and approximates this system by training a sequential autoencoder. We fixed the number of factors in the model to 20 for both the arm and the hand datasets. We then performed PCA on the preprocessed neural activity and kept the components that cumulatively explained 90% of variance in the neural data.

##### Kalman Filter

To predict hand and arm kinematics, we applied the Kalman filter ^28^, commonly used for kinematic decoding ^29,30^. In this approach, kinematic dynamics can be described by a linear relationship between past and future states:

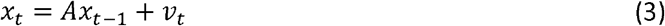

where *x*_*t*_ is a vector of joint angles at time *t*, A is a state transition matrix, and 𝑣_*t*_ is a vector of random numbers drawn from a Gaussian distribution with zero mean and covariance matrix *V*. The kinematics *x*_*t*_ can be also explained in terms of the observed neural activity *z*_*t* − Δ_:

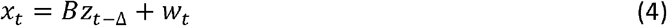

Here, *z*_*t* − Δ_ is a vector of instantaneous firing rates across a population of M1 neurons at time *t* − Δ, *B* is an observation model matrix, and *w*_*t*_ is a random vector drawn from a Gaussian distribution with zero mean and covariance matrix *W*. We tested multiple values of the latency, Δ, and report decoders using the latency that maximized decoder accuracy (150 ms).

We estimated the matrices *A, B, V, W* using linear regression on each training set, and then used those estimates in the Kalman filter update algorithm to infer kinematics of each corresponding test set (see Faragher 2012 and Okorokova et al. 2015 for details) ^31,32^. Briefly, at each time *t*, kinematics were first predicted using the state transition equation (3), then updated with observation information from equation (4). Update of the kinematic prediction was achieved by a weighted average of the two estimates from (3) and (4): the weight of each estimate was inversely proportional to its uncertainty (determined in part by *V* and *W* for the estimates based on *x*_*t-1*_ and *z*_*t-Δ*_, respectively), which changed as a function of time and was thus recomputed for every time step.

To assess decoding performance, we performed 10-fold cross-validation in which we trained the parameters of the filter on a randomly selected 90% of the trials and tested the model using the remaining 10% of trials. Performance was quantified using the average coefficient of determination (*R*^2^) for the held-out trials across test sets. We report performance for each degree of freedom separately (Figure S3) and also the performance averaged across all degrees of freedom (Figure 2).

### Data availability

The data that support the findings of this study are available from the corresponding author upon reasonable request.

### Code availability

The custom analysis code used in this study is available from the corresponding author upon reasonable request.

## Acknowledgments

We would like to thank Sangwook Lee for help with data collection. This work was supported by NINDS grants NS082865, NS101325, and NS096952.

## SUPPLEMENTAL MATERIALS

**Figure S1.**
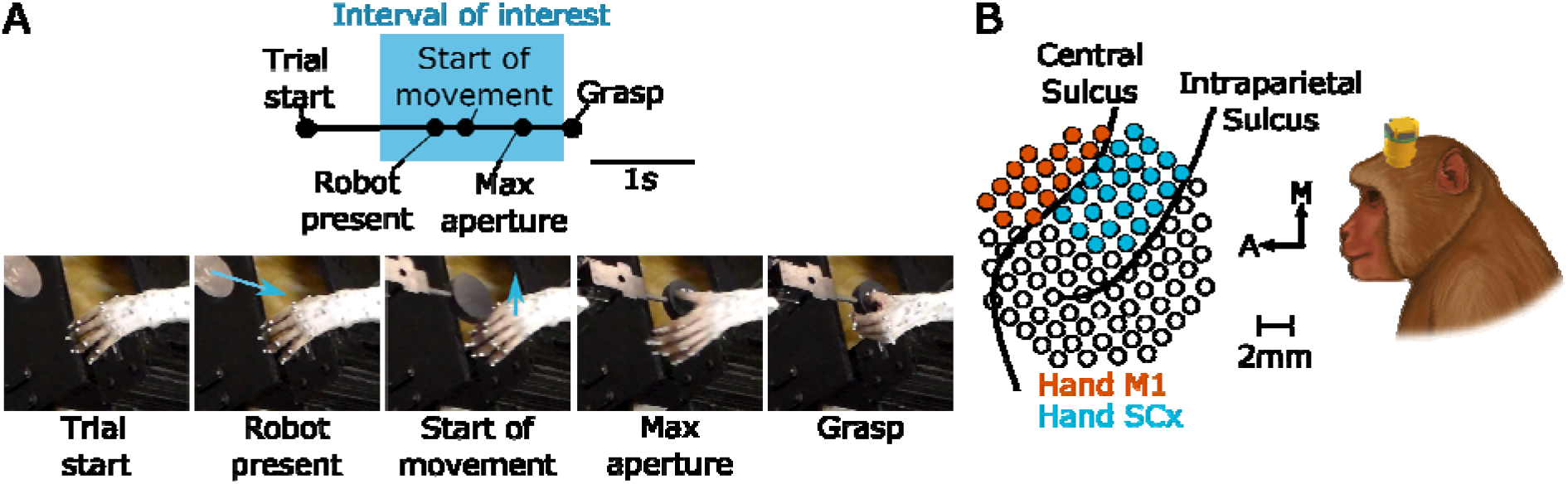
Grasp Task Behavior and Neurophysiology. A| Task Intervals: Start of Movement, Maximum Aperture, and Grasp epochs were manually scored from video. Arrows indicate motion of the robot presenting the object or of the hand. B| Multi-electrode arrays were used to record neuronal activity. The array spanned M1 and SCx, but only M1 units were used for this study except when explicitly noted (i.e., in the tangling analysis).

**Figure S2:**
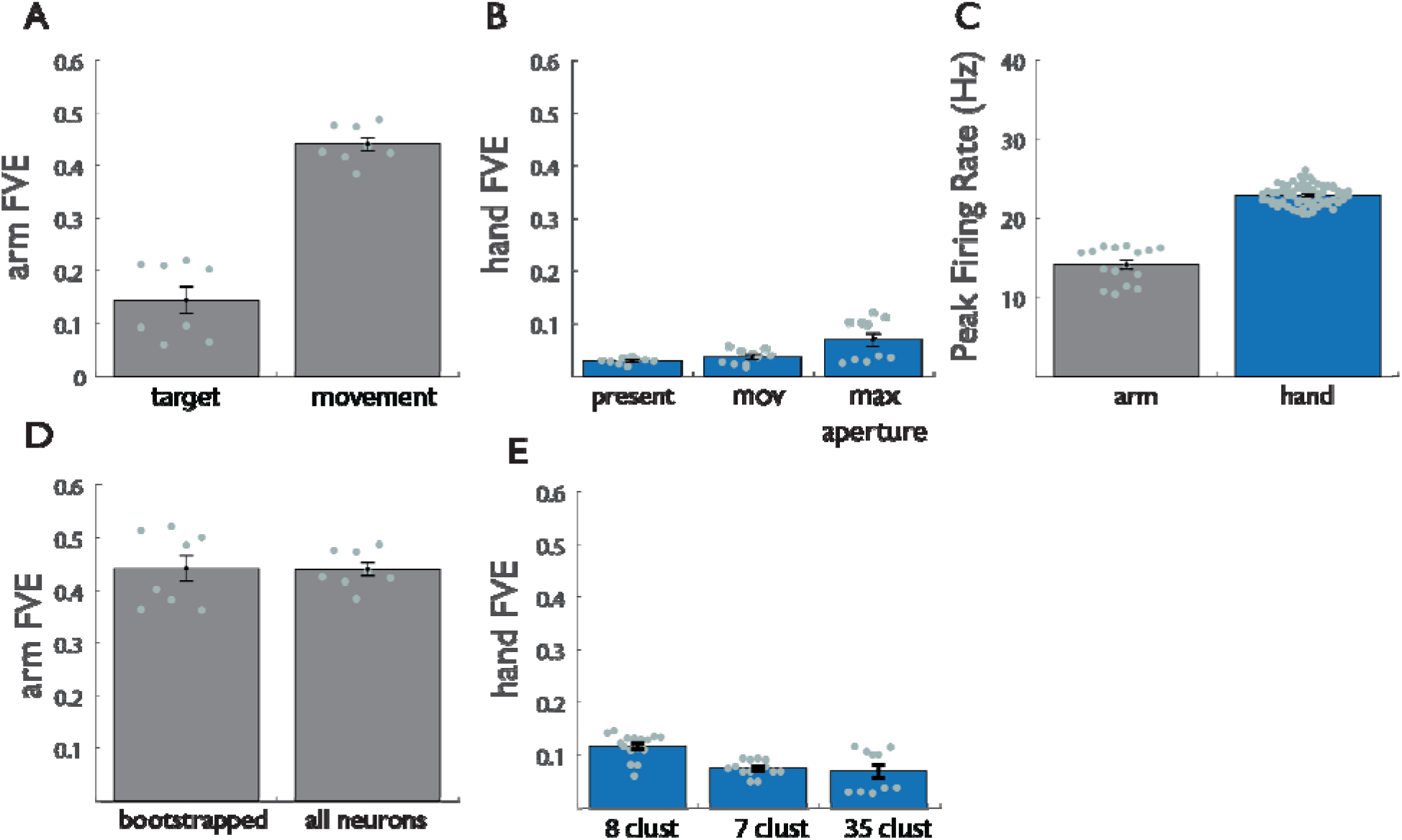
Control analyses for reaching and grasping. A| For reaching: Cross-validated FVE (fraction of variance explained) in the rate of change of neural PCs (d*x*/dt) explained by the optimal linear dynamical system, with data aligned to target presentation (target) or movement onset (movement). B| For grasping: Cross-validated FVE in the rate of change of neural PCs (dx/dt) explained by the optimal linear dynamical system, when the data are aligned to object presentation (present), movement onset (mov), and maximum aperture (max aperture). C| Peak firing rate for arm (gray) and hand (blue) responses. D| Bootstrapped arm responses (55 neurons) vs. full arm dataset. E| Cross-validated fraction of variance explained (FVE) in the rate of change of neural PCs (dx/dt) explained by the optimal linear dynamical system when the objects are clustered into fewer categories for the hand (see methods). Difference between 8 clusters and 35 clusters is significant (*p*=.0008) while difference between 7 clusters and 35 clusters is not significant (*p*=0.57). However, for both clustering methods, difference between hand and arm remains highly significant (8 clusters| *p*=2.5e-18; 7 clusters | *p*=2.08e-19). For all figures, error bars represent standard error of the mean, and data points represent cross-validated results across 2 monkeys.

**Figure S3:**
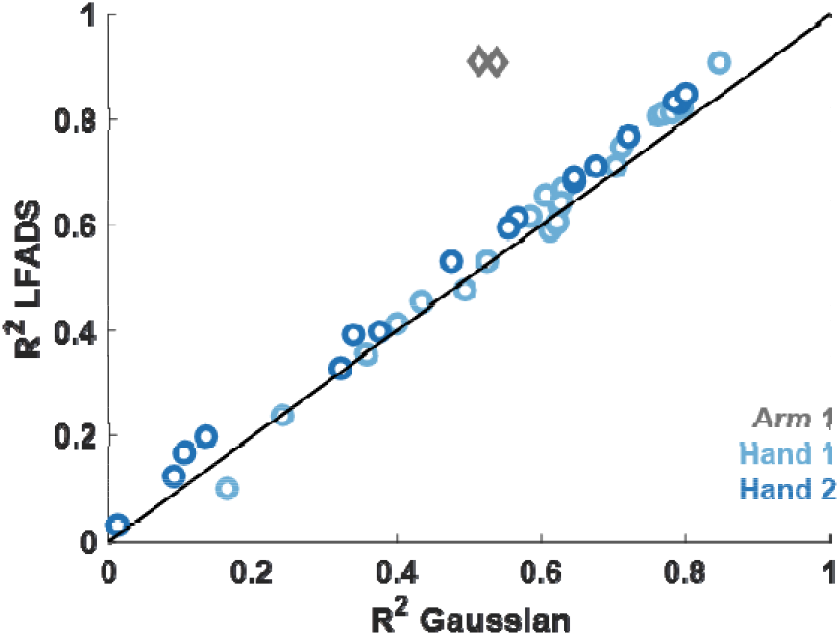
Decoding performance is consistent across joints. Mean performance of separate joints (individual points) derived from decoders with Gaussian smoothing or LFADS preprocessing for 1 arm dataset (grey: N = 44; 2 joints) and 2 hand datasets (light blue: N=44, dark blue: N = 36; 30 joints). For all joints, LFADS leads to substantial improvement in decoding for the arm but not for the hand.

**Figure S4:**
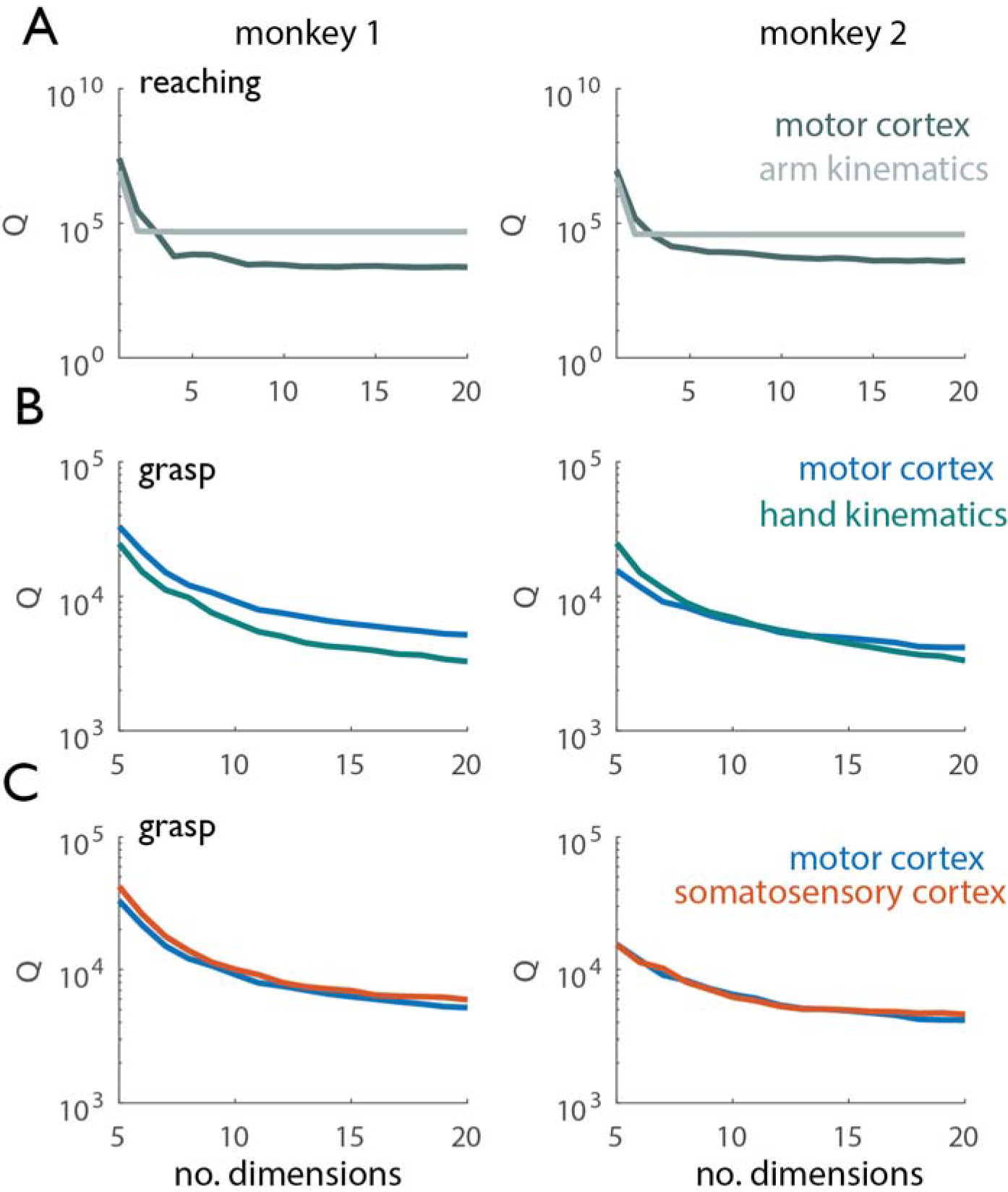
Tangling vs. dimensionality. A| Tangling metric (90^th^ percentile of Q) vs. number of dimensions used to compute Q for reaching. Q values derived from motor cortical responses are shown in dark gray, Q values derived from kinematics are shown in light gray. Arm kinematics exhibit consistently higher tangling than do the corresponding population responses in motor cortex. B| Tangling metric vs. number of dimensions used to compute Q for grasp. Q values derived from motor cortical responses are shown in blue, Q-values derived from hand kinematics are shown in green. When Q has leveled off for the kinematic and neural data (~15 dimensions), neuronal trajectories in motor cortex exhibit higher tangling than do the corresponding hand kinematic trajectories. C| Tangling metric vs. number of dimensions used to compute Q for reaching in motor and somatosensory cortex. Q-values derived from motor cortical responses are shown in blue, those derived from somatosensory responses are shown in orange. Hand motor and somatosensory responses exhibit similar tangling.

## References

1. Churchland, M. M. et al. Neural population dynamics during reaching. Nature 487, 51–56 (2012).

2. Shenoy, K. V, Sahani, M. & Churchland, M. M. Cortical Control of Arm Movements: A Dynamical Systems Perspective. Annu. Rev. Neurosci 36, 337–359 (2013).

3. Russo, A. A. et al. Motor Cortex Embeds Muscle-like Commands in an Untangled Population Response. Neuron 97, 953–966.e8 (2018).

4. Lara, A. H., Elsayed, G. F., Zimnik, A. J., Cunningham, J. P. & Churchland, M. M. Conservation of preparatory neural events in monkey motor cortex regardless of how movement is initiated. Elife 7, e31826 (2018).

5. Hatsopoulos, N. G., Xu, Q. & Amit, Y. Encoding of movement fragments in the motor cortex. J. Neurosci. 27, 5105–5114 (2007).

6. Rouse, A. G. & Schieber, M. H. Condition-Dependent Neural Dimensions Progressively Shift during Reach to Grasp. Cell Rep. 25, 3158–3168.e3 (2018).

7. Remington, E. D., Narain, D., Hosseini, E. A., Correspondence, J. & Jazayeri, M. Flexible Sensorimotor Computations through Rapid Reconfiguration of Cortical Dynamics. Neuron 98, 1005–1019.e5 (2018).

8. Polit, A. & Bizzi, E. Processes controlling arm movements in monkeys. Science 201, 1235–1237 (1978).

9. Brochier, T., Boudreau, M. J., Paré, M. & Smith, A. M. The effects of muscimol inactivation of small regions of motor and somatosensory cortex on independent finger movements and force control in the precision grip. Exp. brain Res. 128, 31–40 (1999).

10. Pandarinath, C. et al. Neural population dynamics in human motor cortex during movements in people with ALS. Elife 4, e07436 (2015).

11. Mochizuki, Y., Mizutani, T., Shimizu, T. & Kawata, A. Proportional neuronal loss between the primary motor and sensory cortex in amyotrophic lateral sclerosis. Neurosci. Lett. 503, 73–75 (2011).

12. Hallett, M., Shahani, B. T. & Young, R. R. EMG analysis of stereotyped voluntary movements in man. J. Neurol. Neurosurg. Psychiatry 38, 1154–1162 (1975).

13. Jones, E. G., Coulter, J. D. & Hendry, S. H. C. lntracortical Connectivity of Architectonic Fields in the Somatic Sensory, Motor and Parietal Cortex of Monkeys. J Comp Neurol 181, 291–347 (1978).

14. Huerta, M. F. & Pons, T. P. Primary motor cortex receives input from area 3a in macaques. Brain Res. 537, 367–371 (1990).

15. Huffman, K. J. & Krubitzer, L. Area 3a: topographic organization and cortical connections in marmoset monkeys. Cereb. Cortex 11, 849–867 (2001).

16. Dotson, N. M., Hoffman, S. J., Goodell, B. & Gray, C. M. A Large-Scale Semi-Chronic Microdrive Recording System for Non-Human Primates. Neuron 96, 769–782 (2017).

17. Anderson, F. C. & Pandy, M. G. Dynamic Optimization of Human Walking. J. Biomech. Eng. 123, 381–390 (2001).

18. Anderson, F. C. & Pandy, M. G. A Dynamic Optimization Solution for Vertical Jumping in Three Dimensions. Comput. Methods Biomech. Biomed. Engin. 2, 201–231 (1999).

19. de Leva, P. Adjustments to Zatsiorsky-Seluyanov’s segment inertia parameters. J. Biomech. 29, 1223–30 (1996).

20. Delp, S. L. et al. An interactive graphics-based model of the lower extremity to study orthopaedic surgical procedures. IEEE Trans. Biomed. Eng. 37, 757–767 (1990).

21. Dempster, W. T. & Gaughran, G. R. L. Properties of body segments based on size and weight. Am. J. Anat. 120, 33–54 (1967).

22. Holzbaur, K. R. S., Murray, W. M. & Delp, S. L. A Model of the Upper Extremity for Simulating Musculoskeletal Surgery and Analyzing Neuromuscular Control. Ann. Biomed. Eng. 33, 829–840 (2005).

23. Yamaguchi, G. T. & Zajac, F. E. A planar model of the knee joint to characterize the knee extensor mechanism. J. Biomech. 22, 1–10 (1989).

24. Delp, S. L. et al. OpenSim: Open-Source Software to Create and Analyze Dynamic Simulations of Movement. IEEE Trans. Biomed. Eng. 54, 1940–1950 (2007).

25. Saleh, M., Takahashi, K., Amit, Y. & Hatsopoulos, N. G. Encoding of coordinated grasp trajectories in primary motor cortex. J. Neurosci. 30, 17079–90 (2010).

26. Goodman, J. M. et al. Postural Representations of the Hand in Primate Sensorimotor Cortex. bioRxiv 566539 (2019). doi:10.1101/566539

27. Pandarinath, C. et al. Inferring single-trial neural population dynamics using sequential auto-encoders. Nat. Methods 15, 805–815 (2018).

28. Kalman, R. E. A New Approach to Linear Filtering and Prediction Problems. Trans. ASME–Journal Basic Eng. 82, 35–45 (1960).

29. Menz, V. K., Schaffelhofer, S. & Scherberger, H. Representation of continuous hand and arm movements in macaque areas M1, F5, and AIP: a comparative decoding study. J. Neural Eng. 12, 056016 (2015).

30. Wu, W. et al. Modeling and Decoding Motor Cortical Activity Using a Switching Kalman Filter. IEEE Trans. Biomed. Eng. 51, 933–942 (2004).

31. Faragher, R. Understanding the Basis of the Kalman Filter Via a Simple and Intuitive Derivation. IEEE Signal Process. Mag. 128–132 (2012).

32. Okorokova, E., Lebedev, M., Linderman, M. & Ossadtchi, A. A dynamical model improves reconstruction of handwriting from multichannel electromyographic recordings. Front. Neurosci. 9, 1–15 (2015).

